# Mitochondrial DNA Copy Number and Incident Atrial Fibrillation

**DOI:** 10.1101/848085

**Authors:** Di Zhao, Traci M. Bartz, Nona Sotoodehnia, Wendy S. Post, Susan R. Heckbert, Alvaro Alonso, Ryan J. Longchamps, Christina A. Castellani, Yun Soo Hong, Jerome I. Rotter, Henry J. Lin, Brian O’Rourke, Nathan Pankratz, John A Lane, Stephanie Y. Yang, Eliseo Guallar, Dan E. Arking

**Author notes:** **Co-Correspondence:** Dan E. Arking, PhD, Johns Hopkins University School of Medicine, 733 N. Broadway, Miller Research Building, Room 459, Baltimore, MD 21205, Eliseo Guallar, MD, DrPH, Welch Center for Prevention, Epidemiology and Clinical Research, Johns Hopkins Bloomberg School of Public Health, 2024 E. Monument Street, Room 2-645, Baltimore, MD 21205.

## Abstract

**Background:** Mechanistic studies suggests that mitochondria DNA (mtDNA) dysfunction may be associated with increased risk of atrial fibrillation (AF). The association between mtDNA copy number (mtDNA-CN) and incident AF in the general population, however, remains unknown.

**Methods:** We conducted prospective analyses of 19,709 participants from the Atherosclerosis Risk in Communities Study (ARIC), the Multi-Ethnic Study of Atherosclerosis (MESA) and the Cardiovascular Health Study (CHS). mtDNA-CN from peripheral blood was calculated from probe intensities on the Affymetrix Genome-Wide Human single nucleotide polymorphisms (SNP) Array 6.0 in ARIC and MESA, and from multiplexed real time quantitative polymerase chain reaction (qPCR) in CHS. Incident AF cases were identified through electrocardiograms, review of hospital discharge codes, Medicare claims, and death certificates.

**Results:** The median follow-up time was 21.4 years in ARIC, 12.9 years in MESA and 11.0 years in CHS, during which 4,021 participants developed incident atrial fibrillation (1,761 in ARIC, 790 in MESA, and 1,470 in CHS). The fully-adjusted pooled hazard ratio for incident atrial fibrillation comparing the 1^st^ to the 5^th^ quintile of mitochondria DNA copy number was 1.13 (1.01, 1.27). The fully-adjusted pooled hazard ratio comparing the 10^th^ vs the 90^th^ percentile of mitochondria DNA copy number was 1.13 (1.04, 1.24). Dose-response spline analysis also showed an inverse association between mitochondria DNA copy number and hazard for atrial fibrillation for all three cohorts. These associations were consistent across subgroups.

**Conclusions:** Mitochondria DNA copy number was inversely associated with the risk of AF independent of traditional cardiovascular risk factors. These findings implicate mitochondria DNA copy number as a novel risk factor for atrial fibrillation. Further research is warranted to understand the underlying mechanisms and to evaluate the role of mitochondria DNA copy number in the management of atrial fibrillation risk.

## Introduction

Atrial fibrillation (AF) is the most common form of clinical cardiac arrhythmia, with rising prevalence and incidence worldwide. [1] The lifetime risk of developing AF ranges from 20% to 37% in Whites and Blacks,[2–4] and it is estimated that the number of adults with AF will double in the US by year 2050, affecting more than 8 million people.[5] AF imposes considerable mortality and morbidity risks related to cardiovascular events and thromboembolism, and is associated with tremendous healthcare costs.[1, 6] The high lifetime risk and adverse consequences of AF highlight the critical need for identifying novel risk markers that may provide insights into AF prevention and treatment.

Mitochondria generate energy for the cell through converting nutrients and oxygen into adenosine triphosphate (ATP).[7] Unlike other organelles, mitochondria have their own circular DNA (mtDNA), which encodes essential genes for oxidative phosphorylation. Each cell contains on average 10^3^ to 10^4^ copies of mtDNA, with variations by cell type and development phase.[8] Mitochondrial DNA copy number (mtDNA-CN) is proportional to transcription of mitochondrial genes and is a marker of mitochondrial dysfunction.[9] Indeed, reduced mtDNA-CN from peripheral blood is associated with adverse cardiovascular disease (CVD) events including heart failure, all-cause mortality, sudden cardiac death, and atherosclerotic CVD,[10–13] as well as with CVD risk factors, including hypertension, diabetes, atherosclerosis and chronic kidney disease.[14–16]

Emerging evidence from mechanistic studies suggests that mtDNA dysfunction may be associated with increased risk of AF through reduced ATP production and elevated reactive oxygen species.[17, 18] The association between mtDNA-CN and incident AF in the general population, however, remains unknown. In the present study, we examined the prospective association between baseline mtDNA-CN and the risk of incident AF among participants from 3 community-based prospective cohort studies: the Atherosclerosis Risk in Communities (ARIC) study, the Multi-Ethnic Study of Atherosclerosis (MESA), and the Cardiovascular Health Study (CHS).

## Methods

### Study population

ARIC is a prospective cohort study of 15,792 men and women 45–64 years of age at baseline (1987–1989).[19] Participants were randomly selected from 4 communities in the US: Forsyth County, NC; Jackson, MS; Minneapolis suburbs, MN; and Washington County, MD. We excluded participants who reported race other than black or white (n=48), blacks from the Minnesota and Maryland centers because the numbers are too small for adequate adjustment or within-community comparisons (n=55), and participants with prevalent CHD (n=667) or prevalent AF at the time of mtDNA-CN measurement (n=292). We further excluded participants whose mtDNA-CN was not measured due to sample or measurement availability (n=4,259), and those missing other covariates (n=322). The final sample included 10,149 participants (**Figure 1**).

**Figure 1.**
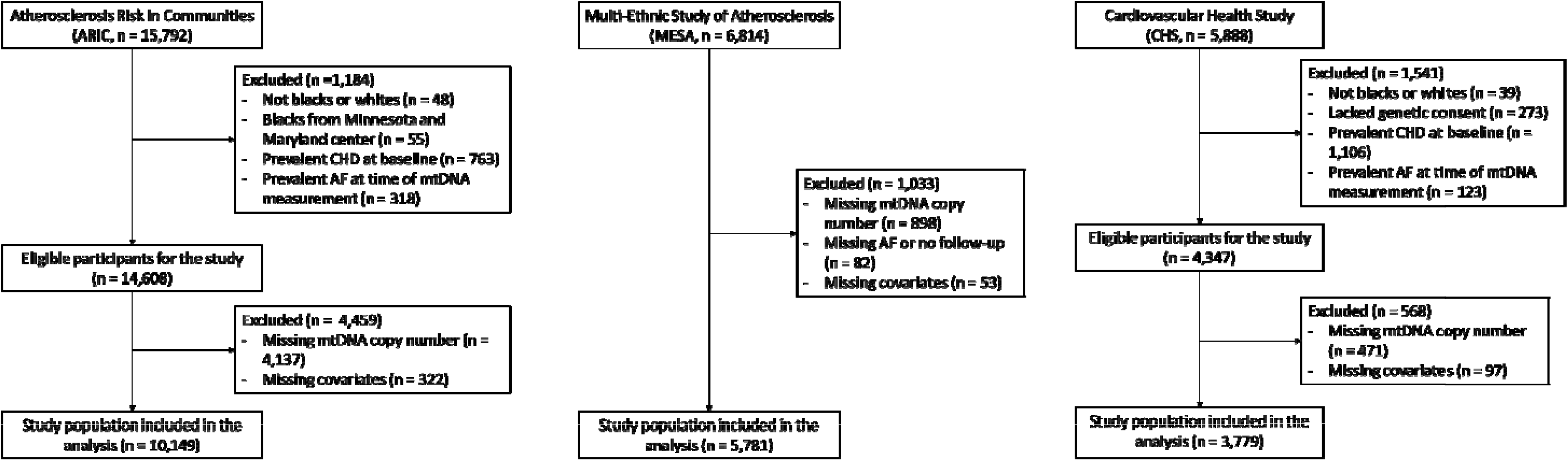
Flowchart of study participants.

MESA is a prospective cohort study of 6,814 men and women aged 45–84 years of age who were free of clinical cardiovascular disease (CVD) at the baseline visit (2000–2002).[20] We excluded participants whose mtDNA-CN was not measured due to sample or technique availability (n=898), who had no information on AF during follow-up (n=82), or who were missing other covariates (n=53). The final study sample included 5,781 participants.

CHS is a prospective cohort study of 5,888 men and women aged 65 years and older.[21] The original cohort of 5201 participants was recruited in 1989-1990 from random samples of Medicare eligibility lists and an additional predominantly African-American cohort of 687 participants was recruited in 1992-1993. We excluded participants who reported race other than black or white (n=39), who lacked genetic consent (n=273), who had prevalent CHD (n=1106) or prevalent AF at the time of mtDNA-CN measurement (n=123). After further excluding participants whose mtDNA-CN was not measured due to sample or technique availability (n=471), and those missing other covariates (n=97), the final study sample included 3,779 participants. Each study obtained approval from their institutional review boards at all centers, and all participants provided written informed consent.

### mtDNA copy number

In ARIC, DNA for mtDNA-CN analysis was collected at visit 1 (1987-1989) for 389 participants (3.8%), at visit 2 (1990-1992) for 8,221 participants (81.0%), at visit 3 (1993-1995) for 1,485 participants (14.6%), and at visit 4 for (1996-1998) for 54 participants (0.5%). In MESA and CHS, DNA for mtDNA-CN analysis was collected at visit 1. The visit of DNA collection was considered as the baseline visit for each participant in the present analysis.

In ARIC and MESA, DNA samples were isolated from buffy coat and genotyped using Affymetrix Genome-Wide Human SNP Arrays 6.0 (the Genvisis software package [www.genvisis.org]).[11, 12, 22] Mitochondrial SNPs were collected across all samples and were signaled with high-quality mitochondrial probes. Unadjusted mtDNA-CN was determined as the median of normalized probe intensity differences across all mitochondrial SNPs. To correct for technical artifacts, batch effects, DNA quality, and starting DNA quantity, we applied surrogate variable analysis (ARIC) and principal component analysis (MESA) to probe intensities of 43,316 Affymetrix autosomal SNPs.[23] We then calculated the residuals of a linear regression model with unadjusted mtDNA-CN as the dependent variable and age, sex, enrollment center, technical covariates, and white blood cell (WBC) count as independent variables in ARIC, and age, sex, collection center, race and principal components as independent variables in MESA. In ARIC, WBC count was missing in 14.9% of participants, and we imputed missing WBC as the study mean. In MESA, rank-based inverse normal transformation was also performed to reduce the impact of outliers.

In CHS, mtDNA-CN was measured using multiplexed real time quantitative polymerase chain reaction (qPCR) utilizing ABI TaqMan chemistry (Applied Biosystems).[12] We calculated residuals using a linear mixed effect model stratified by race, with unadjusted mtDNA-CN as the dependent variable, and age, sex, collection site (fixed effects) and qPCR plate (random effect) as independent variables. The mtDNA-CN residuals were standardized within each study (mean of 0 and standard deviation of 1), and this measure was used as our estimate of mtDNA-CN.

### Atrial fibrillation

In ARIC, AF cases were identified through December 31, 2014 from three sources: electrocardiograms (ECGs) performed during study visits, review of hospital discharge codes, and death certificates.[24] At each study exam, a supine 12-lead resting ECG was performed and transmitted to the ARIC ECG Reading Center (Epidemiological Cardiology Research Center, Wake Forest School of Medicine, Winston Salem, NC) for automatic coding with E Marquette 12-SL program (GE Marquette, Milwaukee, WI). AF or atrial flutter was detected automatically by computer and confirmed by a cardiologist. Hospitalization information during follow-up was obtained through annual follow-up phone calls and surveillance of local hospitals. Trained abstractors collected hospital discharge codes. AF cases detected in the same hospitalization with open cardiac surgery were excluded. The validity of identifying AF from hospital discharge codes has been established in epidemiological studies.[24, 25] The presence of AF was identified if ICD-9-CM codes 427.31 (AF) or 427.32 (atrial flutter) were listed. Finally, AF was identified from death certificates if ICD-9 427.3 or ICD-10 I48 codes were listed as a cause of death. The AF date was determined as the date of the first ECG with AF (4%), the time of first hospital discharge with AF codes (96%), or when AF was listed as a cause of death (0.1%), whichever occurred first.

In MESA, AF cases were identified through December 31, 2014 from three sources: hospital discharge diagnosis codes, Medicare claims data, and study ECGs.[26] Hospitalization information during follow-up was obtained through phone calls every 9-12 months, and medical records and discharge diagnosis were obtained subsequently. Additionally, for participants enrolled in fee-for-service Medicare, AF diagnoses were identified from inpatient, outpatient and physician claims. Study ECGs from visit 5 (2010-2012) were also used to identify incident AF.

In CHS, AF cases were identified through December 31, 2012 from three sources: annual ECGs at each study visit through 1999, discharge diagnoses for all hospitalizations (ICD-9-CM code 427.31 or 427.32), and for those enrolled in fee-for-service Medicare, from inpatient, outpatient or physician claims in Medicare data.[25] The date of AF diagnosis was based on the date of first ECG indicating AF, the time of first hospital discharge with AF codes, or time of the first qualifying outpatient or physician claim, whichever occurred first.

### Other covariates

The measurement of other covariates in the three cohorts has been described previously.[19–21] Age, sex, race/ethnicity, alcohol intake, smoking status, physical activity and medication use were selfreported. Alcohol consumption was categorized into never, former, and current for ARIC and MESA, and into non-current and current for CHS. Body mass index was calculated as weight (kilograms) divided by height (meters) squared. Hypertension was defined as systolic blood pressure ≥ 140 mm Hg, diastolic blood pressure ≥ 90 mm Hg, or current use of anti-hypertension medication. Physical activity was assessed via a modified Baecke Questionnaire in ARIC (scale 1-5), a modified Minnesota Leisure Time Physical Activity Questionnaire in CHS (scale 1-4), and as the total amount of intentional moderate or vigorous exercise performed in a usual week in MESA (MET-min/week). Prevalent heart failure was defined by hospital records, physician diagnosis, or self-reported history of treatment.

Plasma total cholesterol, HDL cholesterol, fasting glucose, and creatinine were measured in each study are previously described.[19–21] N-terminal pro-brain natriuretic peptide (NT-proBNP) was measured at visit 2 and 4 using an electrochemiluminescent immunoassay in ARIC, and at baseline using the Elecsys 2010 analyzer in MESA and CHS. Diabetes was defined as fasting glucose ≥ 126 mg/dL, non-fasting glucose ≥ 200 mg/dL, or use of glycemic control medication. Estimated glomerular filtration rate (eGFR) was calculated using the Chronic Kidney Disease Epidemiology Collaboration (CKD-EPI) equation ARIC and MESA,[27] and Modification of Diet in Renal Disease (CKD-MDRD) equation in CHS.[28] Kidney disease was defined as eGFR < 60 mL/min/1.73 m^2^.

### Statistical analyses

Follow-up started from the baseline visit and continued until development of AF, death, dropout, or through December 31, 2014 in ARIC and MESA or December 31, 2012 in CHS, whichever occurred first. mtDNA-CN was categorized into cohort-specific quintiles. We used a Cox proportional hazards model to estimate hazard ratios (HR) and 95% confidence intervals (CI) for the association between mtDNA-CN and incident AF in each cohort. HRs compared quintiles 1^st^ – 4^th^ with the 5^th^ quintile (reference). Linear trends across quintiles were tested by including a variable containing the median mtDNA-CN level of each quintile in the models. We also modeled mtDNA-CN as a continuous variable and estimated the HR comparing the 10^th^ to the 90^th^ percentile of mtDNA-CN. In addition, to evaluate non-linear dose-response relationships between mtDNA-CN and incident AF, we modeled mtDNA-CN as restricted cubic splines with knots at the 5^th^, 35^th^, 65^th^, and 95^th^ percentiles of its distribution. Finally, we tested for potential interactions by age, sex, race, smoking, alcohol intake, BMI, hypertension, diabetes, and kidney disease.

All analyses were conducted separately in each cohort and cohort-specific HRs were combined using a fixed-effects meta-analysis approach. In each cohort, we used 4 multivariate models with progressive degrees of adjustment. Model 1 was adjusted for age, sex, and race-enrollment center groups. Model 2 was further adjusted for body mass index, height, smoking, alcohol intake, and physical activity. Model 3 was further adjusted for total and HDL cholesterol, cholesterol medication, hypertension, diabetes and prevalent HF. Model 4 was further adjusted for NT-proBNP. Statistical analyses were performed using Stata version 15 for ARIC and MESA study and Stata version 12 for CHS study (StataCorp LP, College Station, Texas). All P values were 2-sided, and statistical significance was declared at P□<□0.05.

## Results

The study included 19,709 participants total (10,149 from ARIC, 5,781 from MESA, and 3,779 from CHS). The mean (standard deviation) age of study participants was 57.2 (5.9), 62.3 (10.3), and 72.2 (5.3) for ARIC, MESA, and CHS, respectively (**Table 1**); 43.5% of participants were men (ARIC 42.4%, MESA 47.8%, and CHS 39.9%) and 69.4% participants were Whites (ARIC 79.0%, MESA 42.6%, and CHS 84.7%). Participants with lower mtDNA-CN were more likely to be current smokers, to have higher NT-proBNP, and to have diabetes and prevalent CKD (**Supplementary Tables 1–3**).

**Table 1.** Baseline characteristics of study participants.

| Characteristic | ARIC | MESA | CHS |
| --- | --- | --- | --- |
| **N** | 10,149 | 5,781 | 3,779 |
| **Age (years)** | 57.2 (5.9) | 62.3 (10.3) | 72.2 (5.3) |
| **Male** | 4,307 (42.4) | 2,762 (47.8) | 1,506 (39.9) |
| **Race** |  |  |  |
| **White** | 8,018 (79.0) | 2,464 (42.6) | 3199 (84.7) |
| **Black** | 2,131 (21.0) | 1,387 (24.0) | 580 (15.3) |
| **Chinese-American** | - | 764 (13.2) | - |
| **Hispanic** | - | 1,166 (20.2) | - |
| **Smoking** |  |  |  |
| **Never** | 4,020 (39.6) | 2,925 (50.6) | 1,766 (46.7) |
| **Former** | 3,814 (37.6) | 2,117 (36.6) | 1,535 (40.6) |
| **Current** | 2,315 (22.8) | 739 (12.8) | 478 (12.6) |
| **Current drinker** | 5,886 (58.0) | 3,221 (55.7) | 1,927 (51.0) |
| **Body mass index (kg/m<sup>2</sup>)** | 28.0 (5.5) | 28.2 (5.5) | 26.7 (4.7) |
| **Total cholesterol (mg/dL)** | 209.8 (39.5) | 194.4 (35.7) | 212.9 (38.6) |
| **HDL cholesterol (mg/dL)** | 50.2 (17.1) | 51.0 (14.9) | 55.5 (15.8) |
| **Triglycerides (mg/dL)** | 115.0 (82.0, 162.0) | 112.0 (78.0, 162.0) | 119.0 (91.0, 162.0) |
| **Systolic BP (mm Hg)** | 122.0 (19.1) | 126.4 (21.7) | 136.3 (21.6) |
| **Diastolic BP (mm Hg)** | 72.2 (10.3) | 71.8 (10.3) | 71.2 (11.2) |
| **NT-proBNP (pg/mL)** | 52.2 (29.0, 91.7) | 56.0 (24.9, 115.3) | 96.8 (51.4, 186.5) |
| **Hypertension** | 3,560 (35.1) | 2,569 (44.4) | 2,134 (56.5) |
| **Diabetes** | 1,413 (13.9) | 700 (12.1) | 510 (13.5) |
| **Prevalent HF** | 395 (3.9) | - | 58 (1.5) |
| **Prevalent CKD** | 192 (1.9) | 776 (13.4) | 721 (19.1) |

Values the Table are mean (SD), number (%), or median (IQR).

The median follow-up time was 21.4 (IQR 14.8–23.3) years in ARIC, 12.9 (9.6–13.6) years in MESA and 11.0 (5.9–17.1) years in CHS. During follow-up, 4,027 participants developed incident AF (1,761 in ARIC, 790 in MESA, and 1,470 in CHS). The HRs for incident AF for the 1^st^ – 4^th^ quintiles of mtDNA-CN compared to the 5^th^ quintile are shown in **Table 2**. In the meta-analysis across the 3 cohorts using the fully adjusted model, the pooled HR for AF comparing the 1^st^ to the 5^th^ quintile of mtDNA-CN was 1.13 (1.01, 1.27) **(Figure 2**). The fully-adjusted pooled HR comparing the 10^th^ vs the 90^th^ percentile of mtDNA-CN was 1.13 (1.04, 1.24). Dose-response spline analysis also showed an inverse association between mtDNA-CN and AF for all three cohorts, with an approximately linear trend (the p-values for non-linear spline terms were 0.20, 0.31 and 0.11 in ARIC, MESA, and CHS, respectively; **Figure 3**).

**Figure 2.**
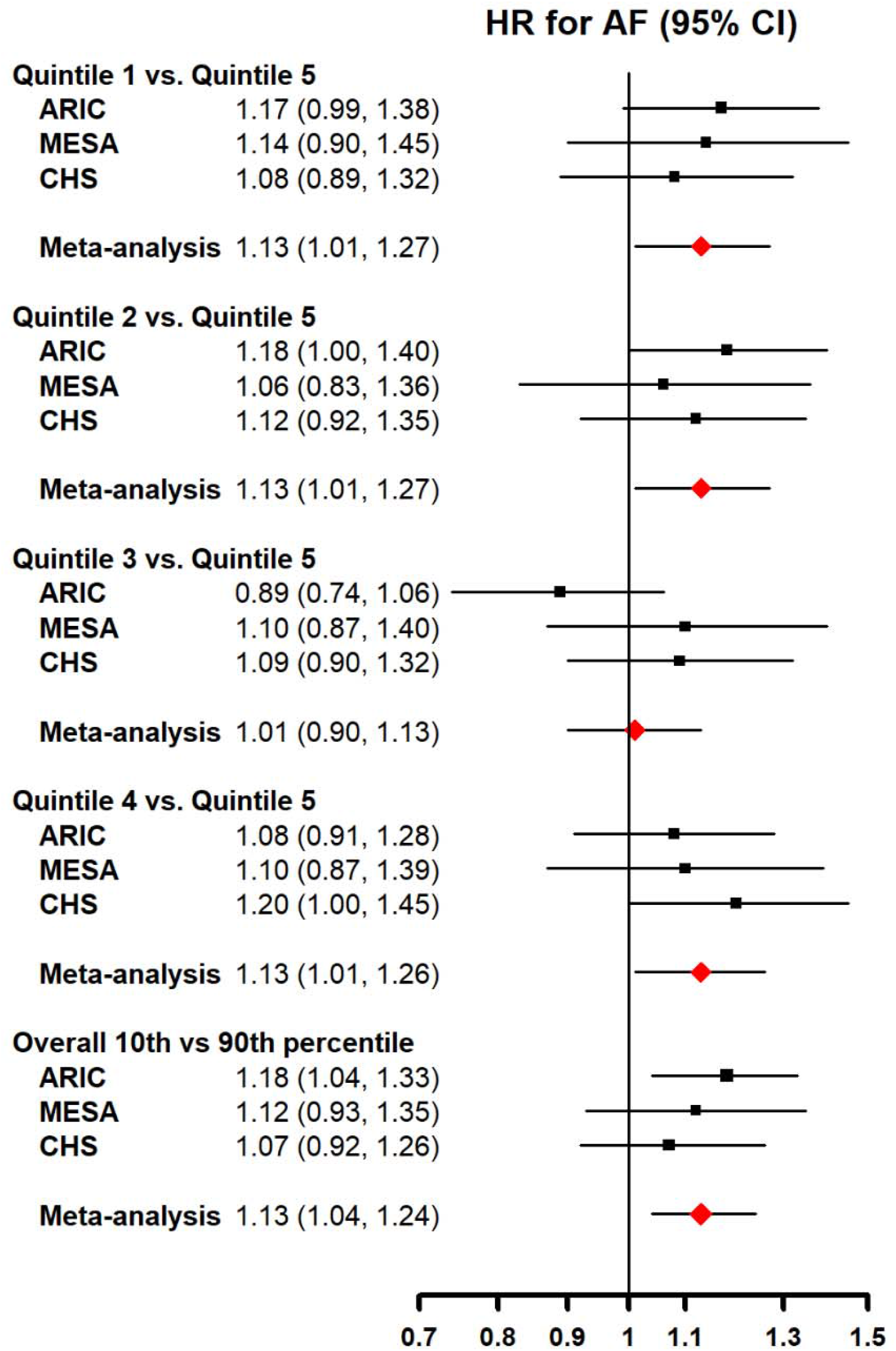
Hazard ratios for incident atrial fibrillation by levels of mtDNA copy number. The figure includes hazard ratios for comparing quintiles 1^st^ – 4^th^ with the 5^th^ quintile (reference) of mtDNA copy number, as well as the hazard ratio for coparing the 10^th^ to the 90^th^ percentile of mtDNA copy number. Models were adjusted for age, sex, race/enrollment center, body mass index, height, smoking, alcohol intake, physical activity, total and HDL cholesterol, cholesterol medication, hypertension, diabetes, prevalent CHD, prevalent heart failure, eGFR and log-transformed NT-proBNP at baseline.

**Figure 3.**
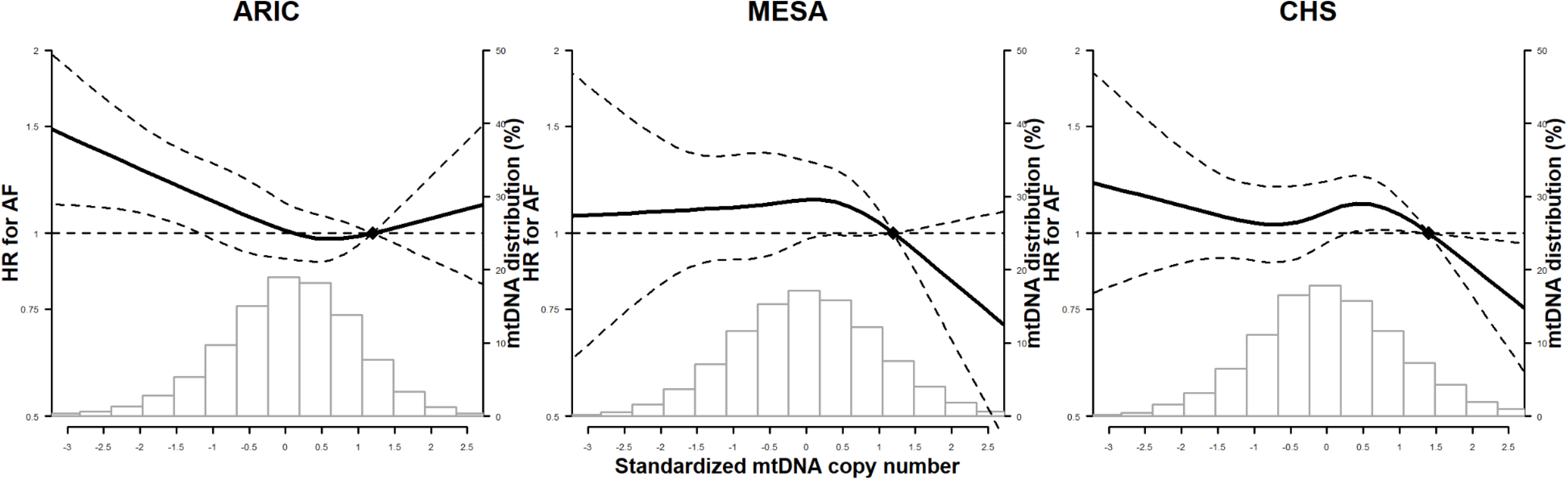
Spline regression analysis of incident atrial fibrillation by levels of mtDNA copy number. The curves represent adjusted hazard ratios (solid line) and their 95% confidence intervals (dashed lines) based on restricted cubic splines of mtDNA copy number with knots at the 5^th^, 35^th^, 65^th^ and 95^th^ percentiles of its distribution. The reference value (diamond dot) was set at the 90^th^ percentile of the distribution. Models were adjusted for age, sex, race/enrollment center, body mass index, height, smoking, alcohol intake, physical activity, total and HDL cholesterol, cholesterol medication, hypertension, diabetes, prevalent CHD, prevalent heart failure, eGFR and log-transformed NT-proBNP at baseline. Histograms represent the frequency distribution of mtDNA copy number at baseline.

**Table 2.**
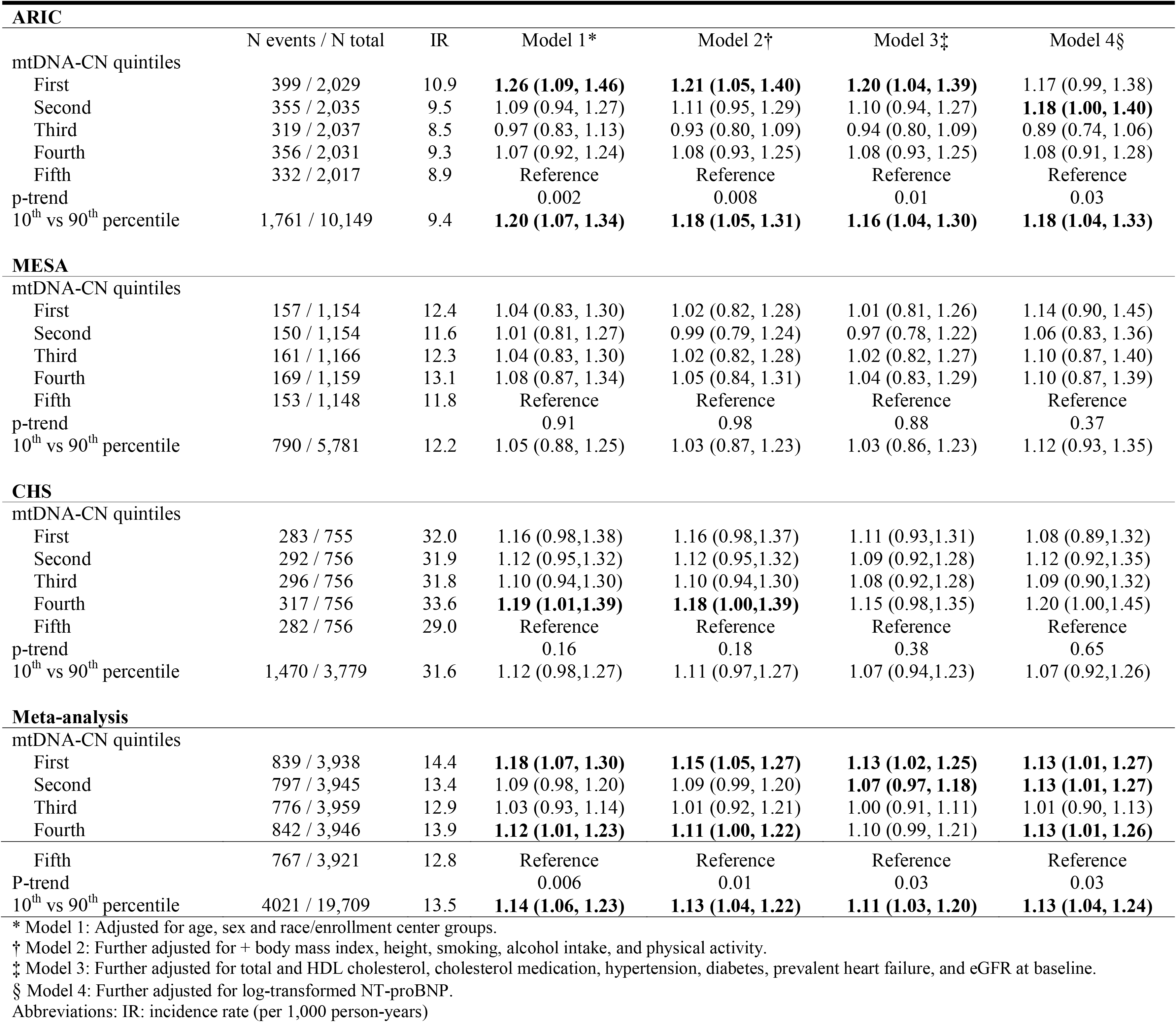
Hazard ratios for incident atrial fibrillation (AF) by quintiles of mtDNA copy number.

| **ARIC** |  |  |  |  |  |  |
| --- | --- | --- | --- | --- | --- | --- |
|  | N events / N total | IR | Model 1* | Model 2† | Model 3‡ | Model 4§ |
| mtDNA-CN quintiles |  |  |  |  |  |  |
| First | 399 / 2,029 | 10.9 | **1.26 (1.09, 1.46)** | **1.21 (1.05, 1.40)** | **1.20 (1.04, 1.39)** | 1.17 (0.99, 1.38) |
| Second | 355 / 2,035 | 9.5 | 1.09 (0.94, 1.27) | 1.11 (0.95, 1.29) | 1.10 (0.94, 1.27) | **1.18 (1.00, 1.40)** |
| Third | 319 / 2,037 | 8.5 | 0.97 (0.83, 1.13) | 0.93 (0.80, 1.09) | 0.94 (0.80, 1.09) | 0.89 (0.74, 1.06) |
| Fourth | 356 / 2,031 | 9.3 | 1.07 (0.92, 1.24) | 1.08 (0.93, 1.25) | 1.08 (0.93, 1.25) | 1.08 (0.91, 1.28) |
| Fifth | 332 / 2,017 | 8.9 | Reference | Reference | Reference | Reference |
| p-trend |  |  | 0.002 | 0.008 | 0.01 | 0.03 |
| 10<sup>th</sup> vs 90<sup>th</sup> percentile | 1,761 / 10,149 | 9.4 | **1.20 (1.07, 1.34)** | **1.18 (1.05, 1.31)** | **1.16 (1.04, 1.30)** | **1.18 (1.04, 1.33)** |
| **MESA** |  |  |  |  |  |  |
| mtDNA-CN quintiles |  |  |  |  |  |  |
| First | 157 / 1,154 | 12.4 | 1.04 (0.83, 1.30) | 1.02 (0.82, 1.28) | 1.01 (0.81, 1.26) | 1.14 (0.90, 1.45) |
| Second | 150 / 1,154 | 11.6 | 1.01 (0.81, 1.27) | 0.99 (0.79, 1.24) | 0.97 (0.78, 1.22) | 1.06 (0.83, 1.36) |
| Third | 161 / 1,166 | 12.3 | 1.04 (0.83, 1.30) | 1.02 (0.82, 1.28) | 1.02 (0.82, 1.27) | 1.10 (0.87, 1.40) |
| Fourth | 169 / 1,159 | 13.1 | 1.08 (0.87, 1.34) | 1.05 (0.84, 1.31) | 1.04 (0.83, 1.29) | 1.10 (0.87, 1.39) |
| Fifth | 153 / 1,148 | 11.8 | Reference | Reference | Reference | Reference |
| p-trend |  |  | 0.91 | 0.98 | 0.88 | 0.37 |
| 10<sup>th</sup> vs 90<sup>th</sup> percentile | 790 / 5,781 | 12.2 | 1.05 (0.88, 1.25) | 1.03 (0.87, 1.23) | 1.03 (0.86, 1.23) | 1.12 (0.93, 1.35) |
| **CHS** |  |  |  |  |  |  |
| mtDNA-CN quintiles |  |  |  |  |  |  |
| First | 283 / 755 | 32.0 | 1.16 (0.98,1.38) | 1.16 (0.98,1.37) | 1.11 (0.93,1.31) | 1.08 (0.89,1.32) |
| Second | 292 / 756 | 31.9 | 1.12 (0.95,1.32) | 1.12 (0.95,1.32) | 1.09 (0.92,1.28) | 1.12 (0.92,1.35) |
| Third | 296 / 756 | 31.8 | 1.10 (0.94,1.30) | 1.10 (0.94,1.30) | 1.08 (0.92,1.28) | 1.09 (0.90,1.32) |
| Fourth | 317 / 756 | 33.6 | **1.19 (1.01,1.39)** | **1.18 (1.00,1.39)** | 1.15 (0.98,1.35) | 1.20 (1.00,1.45) |
| Fifth | 282 / 756 | 29.0 | Reference | Reference | Reference | Reference |
| p-trend |  |  | 0.16 | 0.18 | 0.38 | 0.65 |
| 10<sup>th</sup> vs 90<sup>th</sup> percentile | 1,470 / 3,779 | 31.6 | 1.12 (0.98,1.27) | 1.11 (0.97,1.27) | 1.07 (0.94,1.23) | 1.07 (0.92,1.26) |
| **Meta-analysis** |  |  |  |  |  |  |
| mtDNA-CN quintiles |  |  |  |  |  |  |
| First | 839 / 3,938 | 14.4 | **1.18 (1.07, 1.30)** | **1.15 (1.05, 1.27)** | **1.13 (1.02, 1.25)** | **1.13 (1.01, 1.27)** |
| Second | 797 / 3,945 | 13.4 | 1.09 (0.98, 1.20) | 1.09 (0.99, 1.20) | **1.07 (0.97, 1.18)** | **1.13 (1.01, 1.27)** |
| Third | 776 / 3,959 | 12.9 | 1.03 (0.93, 1.14) | 1.01 (0.92, 1.21) | 1.00 (0.91, 1.11) | 1.01 (0.90, 1.13) |
| Fourth | 842 / 3,946 | 13.9 | **1.12 (1.01, 1.23)** | **1.11 (1.00, 1.22)** | 1.10 (0.99, 1.21) | **1.13 (1.01, 1.26)** |
| Fifth | 767 / 3,921 | 12.8 | Reference | Reference | Reference | Reference |
| P-trend |  |  | 0.006 | 0.01 | 0.03 | 0.03 |
| 10<sup>th</sup> vs 90<sup>th</sup> percentile | 4021 / 19,709 | 13.5 | **1.14 (1.06, 1.23)** | **1.13 (1.04, 1.22)** | **1.11 (1.03, 1.20)** | **1.13 (1.04, 1.24)** |

\* Model 1: Adjusted for age, sex and race/enrollment center groups.
† Model 2: Further adjusted for + body mass index, height, smoking, alcohol intake, and physical activity.
‡ Model 3: Further adjusted for total and HDL cholesterol, cholesterol medication, hypertension, diabetes, prevalent heart failure, and eGFR at baseline.
§ Model 4: Further adjusted for log-transformed NT-proBNP.
Abbreviations: IR: incidence rate (per 1,000 person-years)

In the stratified analysis, there was no evidence of interaction for the associations between mtDNA-CN and AF across all subgroups evaluated (**Supplementary Figure 1**), except that they were stronger in hypertensive than in non-hypertensive participants from ARIC (*P*-interaction=0.03).

## Discussion

In three large population-based prospective cohort studies, mtDNA-CN was inversely associated with the risk of incident AF, independent of traditional risk factors. The association was not statistically different across race and sex groups. This novel association indicates a potential role of mitochondrial dysfunction in atrial arrhythmias, and adds to the pathophysiological evidence from basic science studies supporting a role of mitochondrial mechanisms in the genesis of AF.

Animal models and molecular studies suggest that mitochondrial dysfunction is associated with adverse CVD outcomes and subclinical atherosclerosis,[29, 30] and some of these associations have been confirmed in population-based studies. In ARIC, MESA, and CHS, low levels of mtDNA-CN were associated with increased risk of incident CVD, coronary heart disease, sudden cardiac death, and allcause mortality.[10–12] Low levels of mtDNA-CN were also associated with CVD risk factors, including hypertension, diabetes, and chronic kidney disease.[14–16] Since clinical CVD events and CVD risk factors are also risk factors for AF, the inverse associations between mtDNA-CN and AF could also be mediated through the traditional CVD pathways. As participants in our analysis were free of CHD at baseline and we adjusted for CVD risk factors, our findings suggest that CVD and traditional risk factors do not fully explain the association between mtDNA-CN and incident AF. Furthermore, since low mtDNA-CN levels preceded the development of AF and other CVD events in ARIC, CHS, and MESA, maintaining CVD health may require adequate cell energy mechanisms and mitochondrial function for preserved cardiac contractility, electrical activity, and endothelial function.

AF is triggered by structural and electrophysiological remodeling changes in the atrial myocardium, which in turn adversely affect cardiac function and increase the risk of mortality, stroke, and peripheral embolism.[31] Risk factors of AF include older age, male sex, and the presence of clinical cardiomyopathy, coronary artery disease, and traditional CVD risk factors (hypertension, diabetes, obesity and smoking).[25, 32, 33] Our results suggest that mtDNA-CN may be a novel risk factor for AF. As low levels of mtDNA-CN are a marker for mitochondrial dysfunction and abnormal ATP production, our findings suggest that myocyte electrical activity could be compromised by mitochondrial dysfunction and insufficient energy supply.

The mechanisms underlying the association between mtDNA-CN and AF are unknown, but previous mechanistic research and in-vitro studies provide some leads. Most of the energy for cardiomyocyte electrical activity and cardiac muscle contraction is supplied by mitochondria through the oxidative phosphorylation pathway.[18] The generation of cellular reactive oxygen species (ROS) associated with mitochondrial dysfunction could increase arrhythmia susceptibility by affecting energy-dissipating ion channels and transporters, including the proteins involved in excitation-contraction coupling. Indeed, disrupted intracellular Ca^2+^ homeostasis has been shown to contribute to the pathogenesis of AF.[17, 18, 34] Increased ROS can also impair gap junction regulation and affect voltage-gated sodium potassium channels, which increases electrical heterogeneity and cause early afterdepolarizations.[18]

Apart from ROS, mitochondrial dysfunction impairs ATP synthesis, affecting cardiomyocyte energy metabolism, sarcolemmal and sarcoplasmic ion channel function, intracellular cation homeostasis, and membrane excitability, which are essential in maintaining the electrical activity of cardiac cells.[34] The reduced energy production promotes the opening of sarcolemmal K_ATP_ channels, which has arrhythmogenic effects by conferring shortened action potential duration and excitation wavelengths.[35]

Some limitations of this study need to be considered. Differences in mtDNA-CN measurement techniques, AF identification methods, and covariate assessment methods may contribute to the heterogeneity of the results across the 3 cohorts. Reassuringly, the direction of the association of mtDNA-CN and incident AF was consistent in ARIC, CHS, and MESA, adding weight to the validity of our findings. mtDNA-CN was collected from peripheral blood, and was thus not a direct measurement of mitochondrial function in atrial myocytes. However, mtDNA from peripheral blood has been shown to be strongly correlated with mtDNA from cardiomyocyte (coefficient of correlation >0.5).[13, 36] Therefore, the mtDNA-CN from peripheral blood could be used a marker for myocardial mitochondrial function. Furthermore, mtDNA-CN was measured only at single time-point, and variability due to changes over time was not captured. Finally, we could not evaluate the association between mtDNA-CN with various subtypes of AF. Since AF was obtained primarily based on hospital discharge codes in all 3 studies, the diagnoses were mostly persistent or permanent forms of AF, whereas asymptomatic paroxysmal AF, the most common type of AF, is often undetected. Future studies with repeated measures of mtDNA markers may provide better evaluation of longitudinal changes in mtDNA function and its impact on AF risk.

The strengths of this study include the prospective design, the long follow-up, the rigorous quality control procedures of the individual cohorts, the large sample size, and the heterogeneous composition of the study population, including men and women from multiple race / ethnicity groups and a wide age range from middle-aged adults to elderly participants.

In conclusion, in three prospective community-based cohorts, mtDNA-CN levels in peripheral blood were inversely associated with the risk of AF independent of traditional risk factors. These findings implicate mtDNA-CN as a novel risk factor for AF. Further research is warranted to better understand the underlying mechanisms, to better characterize the dose-response shape of the association, and to evaluate the role of mtDNA-CN in the prevention and management of AF risk.

## Acknowledgements

The authors thank the other investigators, the staff, and the participants of the ARIC, MESA and CHS studies.

## Supplementary materials

### Supplementary figure legend

**Supplementary Figure 1.**
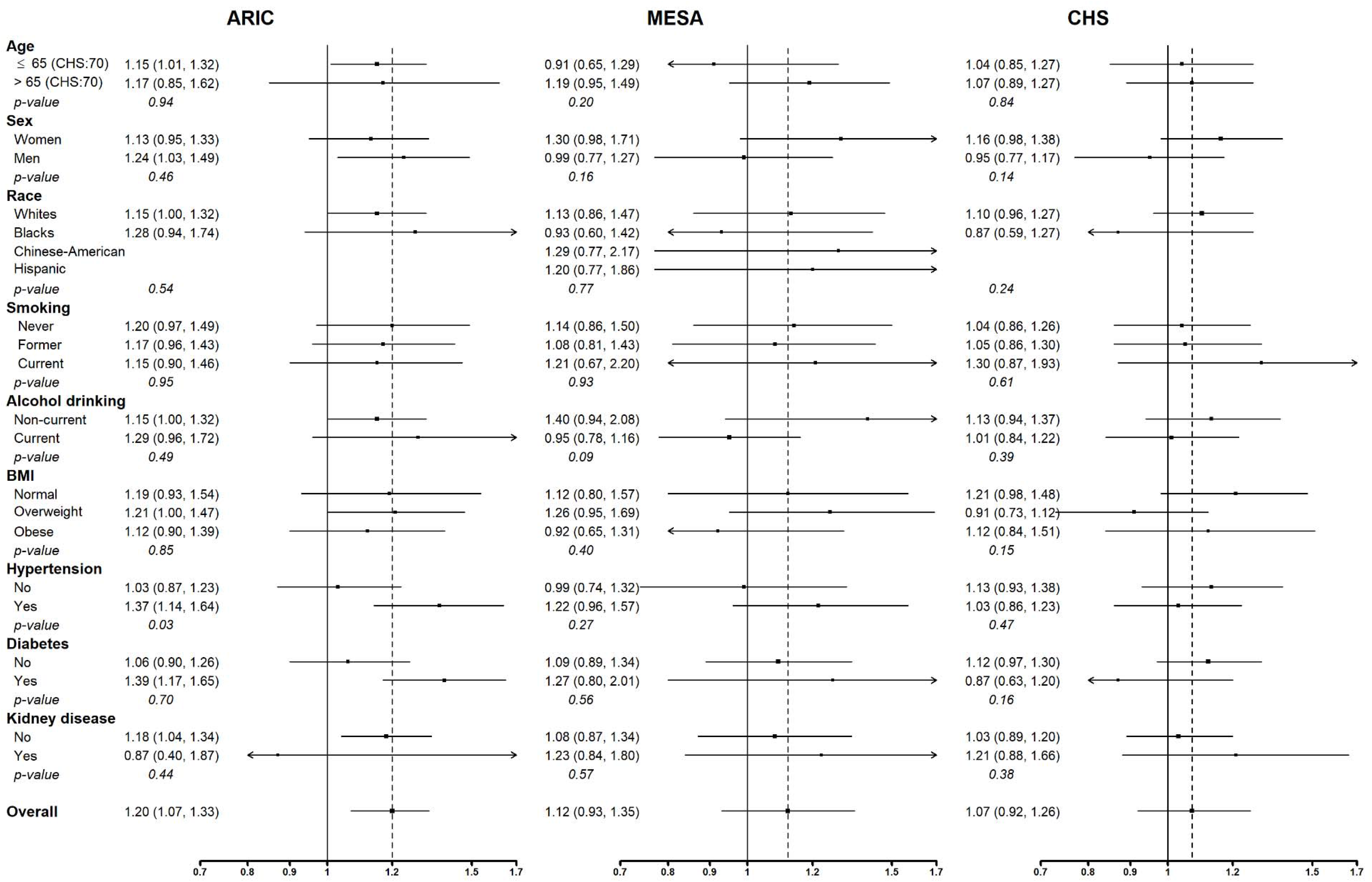
Hazard ratios for incident atrial fibrillation by levels of mtDNA copy number in pre-specified subgroups. Age groups were ≥65 and >65 for ARIC and MESA, ≥70 and >70 for CHS. Hazard ratios are for comparing the 10^th^ to the 90^th^ percentile of mtDNA copy number. Models were adjusted for age, sex, race/enrollment center, body mass index, height, smoking, alcohol intake, physical activity, total and HDL cholesterol, cholesterol medication, hypertension, diabetes, prevalent CHD, prevalent heart failure, eGFR and log-transformed NT-proBNP at baseline.

**Supplementary Table 1.** Baseline characteristics of study participants by mtDNA-CN quintiles in ARIC.

| Characteristic | First | Second | Third | Fourth | Fifth | P-value |
| --- | --- | --- | --- | --- | --- | --- |
| **N** | 2029 | 2035 | 2037 | 2031 | 2017 |  |
| **Age (year)** | 57.2 (6.0) | 57.1 (6.0) | 57.3 (5.9) | 57.0 (5.9) | 57.3 (5.8) | 0.54 |
| **Male** | 849 (41.8) | 861 (42.3) | 863 (42.4) | 847 (41.7) | 887 (44.0) | 0.61 |
| **Race** |  |  |  |  |  | 0.48 |
| White | 1628 (80.2) | 1612 (79.2) | 1606 (78.8) | 1582 (77.9) | 1590 (78.8) |  |
| Black | 401 (19.8) | 423 (20.8) | 431 (21.2) | 449 (22.1) | 427 (21.2) |  |
| **Smoking** |  |  |  |  |  | 0.002 |
| Never | 764 (37.7) | 841 (41.3) | 806 (39.6) | 812 (40.0) | 797 (39.5) |  |
| Former | 733 (36.1) | 727 (35.7) | 775 (38.0) | 784 (38.6) | 795 (39.4) |  |
| Current | 532 (26.2) | 467 (22.9) | 456 (22.4) | 435 (21.4) | 425 (21.1) |  |
| **Current alcohol drinker** | 1125 (55.4) | 1202 (59.1) | 1168 (57.3) | 1228 (60.5) | 1163 (57.7) | 0.02 |
| **Physical activity** | 2.4 (0.8) | 2.5 (0.8) | 2.5 (0.8) | 2.5 (0.8) | 2.5 (0.8) | 0.38 |
| **Body mass index (kg/m<sup>2</sup>)** | 27.9 (5.6) | 27.7 (5.2) | 28.2 (5.6) | 27.8 (5.4) | 28.0 (5.5) | 0.04 |
| **Total cholesterol (mg/dL)** | 209.3 (41.1) | 208.7 (39.2) | 210.0 (37.9) | 209.8 (39.0) | 211.1 (40.3) | 0.39 |
| **HDL cholesterol (mg/dL)** | 48.9 (16.8) | 50.1 (17.2) | 50.3 (16.6) | 51.5 (17.8) | 50.1 (17.1) | <0.001 |
| **Triglycerides (mg/dL)** | 116.0 (83.0, 162.0) | 115.0 (82.0, 161.0) | 116.0 (82.0, 165.0) | 109.0 (81.0, 158.0) | 118.0 (85.0, 162.0) | 0.01 |
| **Systolic BP (mm Hg)** | 122.7 (19.6) | 122.0 (19.7) | 122.1 (18.8) | 121.5 (18.8) | 121.7 (18.5) | 0.33 |
| **Diastolic BP (mm Hg)** | 72.3 (10.5) | 72.2 (10.4) | 72.1 (10.1) | 72.3 (9.9) | 72.3 (10.5) | 0.97 |
| **NT-proBNP (pg/mL)** | 56.5 (31.5, 102.0) | 51.3 (29.4, 92.8) | 52.3 (28.0, 90.9) | 50.5 (28.7, 86.9) | 50.9 (27.6, 86.2) | <0.001 |
| **Hypertension** | 748 (36.9) | 716 (35.2) | 695 (34.1) | 700 (34.5) | 701 (34.8) | 0.39 |
| **Diabetes** | 336 (16.6) | 296 (14.5) | 269 (13.2) | 245 (12.1) | 267 (13.2) | <0.001 |
| **Prevalent HF** | 73 (3.6) | 73 (3.6) | 80 (3.9) | 76 (3.7) | 93 (4.6) | 0.42 |
| **Prevalent CKD** | 53 (2.6) | 38 (1.9) | 34 (1.7) | 33 (1.6) | 34 (1.7) | 0.11 |

Values the Table are mean (SD), number (%), or median (IQR)

**Supplementary table 2.** Baseline characteristics of study participants by mtDNA-CN quintiles in MESA.

| Characteristic | First | Second | Third | Fourth | Fifth | P-value |
| --- | --- | --- | --- | --- | --- | --- |
| **N** | 1,154 | 1,154 | 1,166 | 1,159 | 1,148 |  |
| **Age (year)** | 62.5 (10.4) | 62.0 (10.3) | 62.2 (10.1) | 62.5 (10.4) | 62.4 (10.1) | 0.78 |
| **Male** | 540 (46.8) | 556 (48.2) | 573 (49.1) | 547 (47.2) | 546 (47.6) | 0.81 |
| **Race / enrollment center** |  |  |  |  |  | 0.10 |
| White | 492 (42.6) | 511 (44.3) | 486 (41.7) | 493 (42.5) | 482 (42.0) |  |
| Black | 140 (12.1) | 147 (12.7) | 167 (14.3) | 172 (14.8) | 138 (12.0) |  |
| Chinese-American | 303 (26.3) | 261 (22.6) | 254 (21.8) | 270 (23.3) | 299 (26.0) |  |
| Hispanic | 219 (19.0) | 235 (20.4) | 259 (22.2) | 224 (19.3) | 229 (19.9) |  |
| **Smoking** |  |  |  |  |  | 0.42 |
| Never | 585 (50.7) | 595 (51.6) | 594 (50.9) | 582 (50.2) | 569 (49.6) |  |
| Former | 410 (35.5) | 401 (34.7) | 424 (36.4) | 432 (37.3) | 450 (39.2) |  |
| Current | 159 (13.8) | 158 (13.7) | 148 (12.7) | 145 (12.5) | 129 (11.2) |  |
| **Current alcohol drinker** | 649 (56.2) | 642 (55.6) | 646 (55.4) | 637 (55.0) | 647 (56.4) | 0.96 |
| **Physical activity** | 5638.3 (6338.1) | 5510.9 (5500.7) | 6056.2 (6305.4) | 5404.8 (5591.3) | 5677.2 (5381.9) | 0.04 |
| **Body mass index (kg/m<sup>2</sup>)** | 28.4 (5.8) | 28.2 (5.4) | 28.2 (5.5) | 28.2 (5.4) | 27.9 (5.2) | 0.2 |
| **Total cholesterol (mg/dL)** | 194.5 (36.4) | 193.7 (37.0) | 195.0 (35.5) | 193.9 (33.1) | 195.0 (36.3) | 0.84 |
| **HDL cholesterol (mg/dL)** | 50.9 (14.6) | 50.4 (15.0) | 50.0 (14.1) | 51.4 (15.2) | 52.2 (15.3) | 0.004 |
| **Triglycerides (mg/dL)** | 114.0 (79.0, 166.0) | 113.5 (79.0, 162.0) | 116.5 (79.0, 170.0) | 110.0 (78.0, 159.0) | 106.0 (77.5, 156.0) | 0.08 |
| **Systolic BP (mm Hg)** | 127.8 (20.8) | 127.0 (21.8) | 125.9 (21.3) | 126.2 (22.1) | 125.0 (22.3) | 0.02 |
| **Diastolic BP (mm Hg)** | 72.1 (10.3) | 72.4 (10.2) | 71.7 (10.3) | 71.3 (10.4) | 71.3 (10.1) | 0.03 |
| **NT-proBNP (pg/mL)** | 57.3 (23.8 - 116.3) | 52.4 (25.2 - 110.0) | 51.7 (23.3 - 108.5) | 59.1 (26.8 - 120.9) | 59.8 (26.3 - 120.9) | 0.14 |
| **Hypertension** | 551 (47.7) | 518 (44.9) | 504 (43.2) | 510 (44.0) | 486 (42.3) | 0.09 |
| **Diabetes** | 159 (13.8) | 147 (12.7) | 138 (11.8) | 135 (11.6) | 121 (10.5) | 0.17 |
| **Prevalent CKD** | 185 (16.0) | 150 (13.0) | 162 (13.9) | 139 (12.0) | 140 (12.2) | 0.03 |

Values the Table are mean (SD), number (%), or median (IQR)

**Supplementary table 3.** Baseline characteristics of study participants by mtDNA-CN quintiles in CHS.

| Characteristic | First | Second | Third | Fourth | Fifth | p-value |
| --- | --- | --- | --- | --- | --- | --- |
| **N** | 755 | 756 | 756 | 756 | 756 |  |
| **Age (year)** | 71.9 (5.2) | 72.3 (5.3) | 72.2 (5.3) | 72.5 (5.4) | 72.1 (5.2) | 0.20 |
| **Male gender** | 290 (38.4) | 302 (39.9) | 325 (43.0) | 280 (37.0) | 309 (40.9) | 0.16 |
| **Race** |  |  |  |  |  |  |
| White | 638 (84.5) | 642 (84.9) | 651 (86.1) | 632 (83.6) | 636 (84.1) | 0.71 |
| Black | 117 (15.5) | 114 (15.1) | 105 (13.9) | 124 (16.4) | 120 (15.9) |  |
| **Smoking** |  |  |  |  |  | 0.51 |
| Never | 348 (46.1) | 360 (47.6) | 359 (47.5) | 350 (46.3) | 349 (46.2) |  |
| Former | 298 (39.5) | 294 (38.9) | 303 (40.1) | 314 (41.5) | 326 (43.1) |  |
| Current | 109 (14.4) | 102 (13.5) | 94 (12.4) | 92 (12.2) | 81 (10.7) |  |
| **Current alcohol drinker** | 344 (45.6) | 379 (50.1) | 389 (51.5) | 401 (53.0) | 414 (54.8) | 0.005 |
| **Physical activity** | 2.2 (1.0) | 2.2 (1.0) | 2.3 (1.0) | 2.3 (1.0) | 2.4 (1.0) | 0.001 |
| **Body mass index (kg/m<sup>2</sup>)** | 26.7 (4.9) | 26.5 (4.9) | 26.7 (4.5) | 26.7 (4.7) | 26.7 (4.6) | 0.94 |
| **Total cholesterol (mg/dL)** | 210.6 (39.0) | 211.7 (37.4) | 213.3 (38.0) | 213.7 (39.8) | 215.1 (38.8) | 0.17 |
| **HDL cholesterol (mg/dL)** | 54.4 (15.5) | 56.0 (16.5) | 55.2 (15.6) | 55.7 (15.4) | 56.1 (16.0) | 0.23 |
| **Triglycerides (mg/dL)** | 122.0 (93.0,<br>170.0) | 116.0 (93.5,<br>155.0) | 117.0 (90.0,<br>159.5) | 119.0 (90.0,<br>164.5) | 120.0 (90.0,<br>160.0) | 0.45 |
| **Systolic BP (mm Hg)** | 135.9 (21.8) | 136.8 (22.2) | 137.2 (21.6) | 136.1 (21.1) | 135.4 (21.4) | 0.48 |
| **Diastolic BP (mm Hg)** | 70.6 (11.3) | 71.5 (11.9) | 72.2 (11.1) | 70.8 (11.0) | 70.8 (10.7) | 0.04 |
| **NT-proBNP (pg/mL)** | 102.0 (52.5 ,193.8) | 110.6 (59.7 ,203.8) | 96.2 (51.5 ,185.0) | 95.0 (50.5 ,184.8) | 86.6 (45.7 ,156.6) | 0.001 |
| **Hypertension** | 424 (56.2) | 433 (57.3) | 442 (58.5) | 432 (57.1) | 403 (53.3) | 0.32 |
| **Diabetes** | 120 (15.9) | 106 (14.0) | 79 (10.4) | 102 (13.5) | 103 (13.6) | 0.04 |
| **Prevalent HF** | 18 (2.4) | 12 (1.6) | 8 (1.1) | 14 (1.9) | 6 (0.8) | 0.09 |
| **Prevalent CKD** | 187 (24.8) | 148 (19.6) | 161 (21.3) | 121 (16.0) | 104 (13.8) | <0.001 |

Values the Table are mean (SD), number (%), or median (IQR)

## References

1. Fuster V, Ryden LE, Cannom DS, Crijns HJ, Curtis AB, Ellenbogen KA, Halperin JL, Kay GN, Le Huezey JY, Lowe JE et al: 2011 ACCF/AHA/HRS focused updates incorporated into the ACC/AHA/ESC 2006 guidelines for the management of patients with atrial fibrillation: a report of the American College of Cardiology Foundation/American Heart Association Task Force on practice guidelines. Circulation 2011, 123(10):e269–367.

2. Lloyd-Jones DM, Wang TJ, Leip EP, Larson MG, Levy D, Vasan RS, D’Agostino RB, Massaro JM, Beiser A, Wolf PA et al: Lifetime risk for development of atrial fibrillation: the Framingham Heart Study. Circulation 2004, 110(9):1042–1046.

3. Weng LC, Preis SR, Hulme OL, Larson MG, Choi SH, Wang B, Trinquart L, McManus DD, Staerk L, Lin H et al: Genetic Predisposition, Clinical Risk Factor Burden, and Lifetime Risk of Atrial Fibrillation. Circulation 2018, l37(lO):1027–1038.

4. Mou L, Norby FL, Chen LY, O’Neal WT, Lewis TT, Loehr LR, Soliman EZ, Alonso A: Lifetime Risk of Atrial Fibrillation by Race and Socioeconomic Status: ARIC Study (Atherosclerosis Risk in Communities). Circ Arrhythm Electrophysiol 2018, 11(7):e006350.

5. Colilla S, Crow A, Petkun W, Singer DE, Simon T, Liu X: Estimates of current and future incidence and prevalence of atrial fibrillation in the U.S. adult population. Am J Cardiol 2013, 112(8):1142–1147.

6. Chugh SS, Havmoeller R, Narayanan K, Singh D, Rienstra M, Benjamin EJ, Gillum RF, Kim YH, McAnulty JH, Jr., Zheng ZJ et al: Worldwide epidemiology of atrial fibrillation: a Global Burden of Disease 2010 Study. Circulation 2014, 129(8):837–847.

7. Friedman JR, Nunnari J: Mitochondrial form and function. Nature 2014, 505(7483):335–343.

8. Rooney JP, Ryde IT, Sanders LH, Howlett EH, Colton MD, Germ KE, Mayer GD, Greenamyre JT, Meyer JN: PCR based determination of mitochondrial DNA copy number in multiple species. Methods Mol Biol 2015, 1241:23–38.

9. Malik AN, Czajka A: Is mitochondrial DNA content a potential biomarker of mitochondrial dysfunction? Mitochondrion 2013, lЗ(5):481–492.

10. Zhang Y, Guallar E, Ashar FN, Longchamps RJ, Castellani CA, Lane J, Grove ML, Coresh J, Sotoodehnia N, Ilkhanoff L et al: Association between mitochondrial DNA copy number and sudden cardiac death: findings from the Atherosclerosis Risk in Communities study (ARIC). Eur Heart J 2017, 38(46):3443–3448.

11. Ashar FN, Zhang Y, Longchamps RJ, Lane J, Moes A, Grove ML, Mychaleckyj JC, Taylor KD, Coresh J, Rotter JI et al: Association of Mitochondrial DNA Copy Number With Cardiovascular Disease. JAMA Cardiol 2017, 2(11):1247–1255.

12. Ashar FN, Moes A, Moore AZ, Grove ML, Chaves PH, Coresh J, Newman AB, Matteini AM, Bandeen-Roche K, Boerwinkle E et al: Association of mitochondrial DNA levels with frailty and all-cause mortality. J Mol Med (Berl) 2015, 93(2):177–186.

13. Huang J, Tan L, Shen R, Zhang L, Zuo H, Wang DW: Decreased Peripheral Mitochondrial DNA Copy Number is Associated with the Risk of Heart Failure and Long-term Outcomes. Medicine (Baltimore) 2016, 95(15):e3323.

14. Tin A, Grams ME, Ashar FN, Lane JA, Rosenberg AZ, Grove ML, Boerwinkle E, Selvin E, Coresh J, Pankratz N et al: Association between Mitochondrial DNA Copy Number in Peripheral Blood and Incident CKD in the Atherosclerosis Risk in Communities Study. J Am Soc Nephrol 2016, 27(8):2467–2473.

15. Lee HK, Song JH, Shin CS, Park DJ, Park KS, Lee KU, Koh CS: Decreased mitochondrial DNA content in peripheral blood precedes the development of non-insulin-dependent diabetes mellitus. Diabetes Res Clin Pract 1998, 42(3):161–167.

16. Tang X, Luo YX, Chen HZ, Liu DP: Mitochondria, endothelial cell function, and vascular diseases. Front Physiol 2014, 5:175.

17. Xie W, Santulli G, Reiken SR, Yuan Q, Osborne BW, Chen BX, Marks AR: Mitochondrial oxidative stress promotes atrial fibrillation. Sci Rep 2015, 5:11427.

18. Yang KC, Bonini MG, Dudley SC, Jr.: Mitochondria and arrhythmias. Free Rodic Biol Med 2014, 71:351–361.

19. The Atherosclerosis Risk in Communities (ARIC) Study: design and objectives. The ARIC investigators. Am J Epidemiol 1989, 129(4):687–702.

20. Bild DE, Bluemke DA, Burke GL, Detrano R, Diez Roux AV, Folsom AR, Greenland P, Jacob DR, Jr., Kronmal R, Liu K eř al: Multi-Ethnic Study of Atherosclerosis: objectives and design. Am J Epidemiol 2002, 156(9):871–881.

21. Fried LP, Borhani NO, Enright P, Furberg CD, Gardin JM, Kronmal RA, Kuller LH, Manolio TA, Mittelmark MB, Newman A et al: The Cardiovascular Health Study: design and rationale. Ann Epidemiol 1991, 1(3):263–276.

22. Longchamps R, Castellani C, Newcomb C, Sumpter J, Lane J, Grove M, Guallar E, Pankratz N, Taylor K, Rotter J et al: Evaluation of mitochondrial DNA copy number estimation techniques. 2019.

23. Leek JT, Johnson WE, Parker HS, Jaffe AE, Storey JD: The sva package for removing batch effects and other unwanted variation in high-throughput experiments. Bioinformotics 2012, 28(6):882–883.

24. Alonso A, Agarwal SK, Soliman EZ, Ambrose M, Chamberlain AM, Prineas RJ, Folsom AR: Incidence of atrial fibrillation in whites and African-Americans: the Atherosclerosis Risk in Communities (ARIC) study. Am Heart J 2009, 158(1):11 1–117.

25. Psaty BM, Manolio TA, Kuller LH, Kronmal RA, Cushman M, Fried LP, White R, Furberg CD, Rautaharju PM: Incidence of and risk factors for atrial fibrillation in older adults. Circulation 1997, 96(7):2455–2461.

26. Heckbert SR, Wiggins KL, Blackshear C, Yang Y, Ding J, Liu J, McKnight B, Alonso A, Austin TR, Benjamin EJ et al: Pericardial fat volume and incident atrial fibrillation in the Multi-Ethnic Study of Atherosclerosis and Jackson Heart Study. Obesity (Silver Spring) 2017, 25(6):1 115–1121.

27. Levey AS, Stevens LA, Schmid CH, Zhang YL, Castro AF, 3rd, Feldman HI, Kusek JW, Eggers P, Van Lente F, Greene T et al: A new equation to estimate glomerular filtration rate. Ann Intern Med 2009, 150(9):604–612.

28. Levey AS, Stevens LA, Schmid CH, Zhang YL, Castro AF, 3rd, Feldman HI, Kusek JW, Eggers P, Van Lente F, Greene T et al: A new equation to estimate glomerular filtration rate. Ann Intern Med 2009, 150(9):604–612.

29. Stockigt F, Beiert T, Knappe V, Baris OR, Wiesner RJ, Clemen CS, Nickenig G, Andrie RP, Schrickel JW: Aging-related mitochondrial dysfunction facilitates the occurrence of serious arrhythmia after myocardial infarction. Biochem Biophys Res Commun 2017, 493(1):604–610.

30. Ballinger SW: Mitochondrial dysfunction in cardiovascular disease. Free Radio Biol Med 2005, 38(10):1278–1295.

31. Chugh SS, Blackshear JL, Shen WK, Hammill SC, Gersh BJ: Epidemiology and natural history of atrial fibrillation: clinical implications. J Am Coll Cardiol 2001, 37(2):371–378.

32. Benjamin EJ, Levy D, Vaziri SM, D’Agostino RB, Belanger AJ, Wolf PA: Independent risk factors for atrial fibrillation in a population-based cohort. The Framingham Heart Study. Jama 1994, 271(11):840–844.

33. Magnani JW, Rienstra M, Lin H, Sinner MF, Lubitz SA, McManus DD, Dupuis J, Ellinor PT, Benjamin EJ: Atrial fibrillation: current knowledge and future directions in epidemiology and genomics. Circulation 2011, 124(18):1982–1993.

34. Montaigne D, Marechal X, Lefebvre P, Modine T, Fayad G, Dehondt H, Hurt C, Coisne A, Koussa M, Remy-Jouet I et al: Mitochondrial dysfunction as an arrhythmogenic substrate: a translational proof-of-concept study in patients with metabolic syndrome in whom post-operative atrial fibrillation develops. J Am Coll Cardiol 2013, 62(16):1466–1473.

35. Brown DA, O’Rourke B: Cardiac mitochondria and arrhythmias. Cardiovasc Res 2010, 88(2):241–249.

36. Knez J, Lakota K, Bozic N, Okrajsek R, Cauwenberghs N, Thijs L, Knezevic I, Vrtovec B, Tomsic M, Cucnik S et al: Correlation Between Mitochondrial DNA Content Measured in Myocardium and Peripheral Blood of Patients with Non-Ischemic Heart Failure. Genet Test Mol Biomarkers 2017, 21(12):736–741.

