## Supplementary figures and images for "Mitochondrial DNA Copy Number and Incident Atrial Fibrillation"

### Supplemental Figure 1

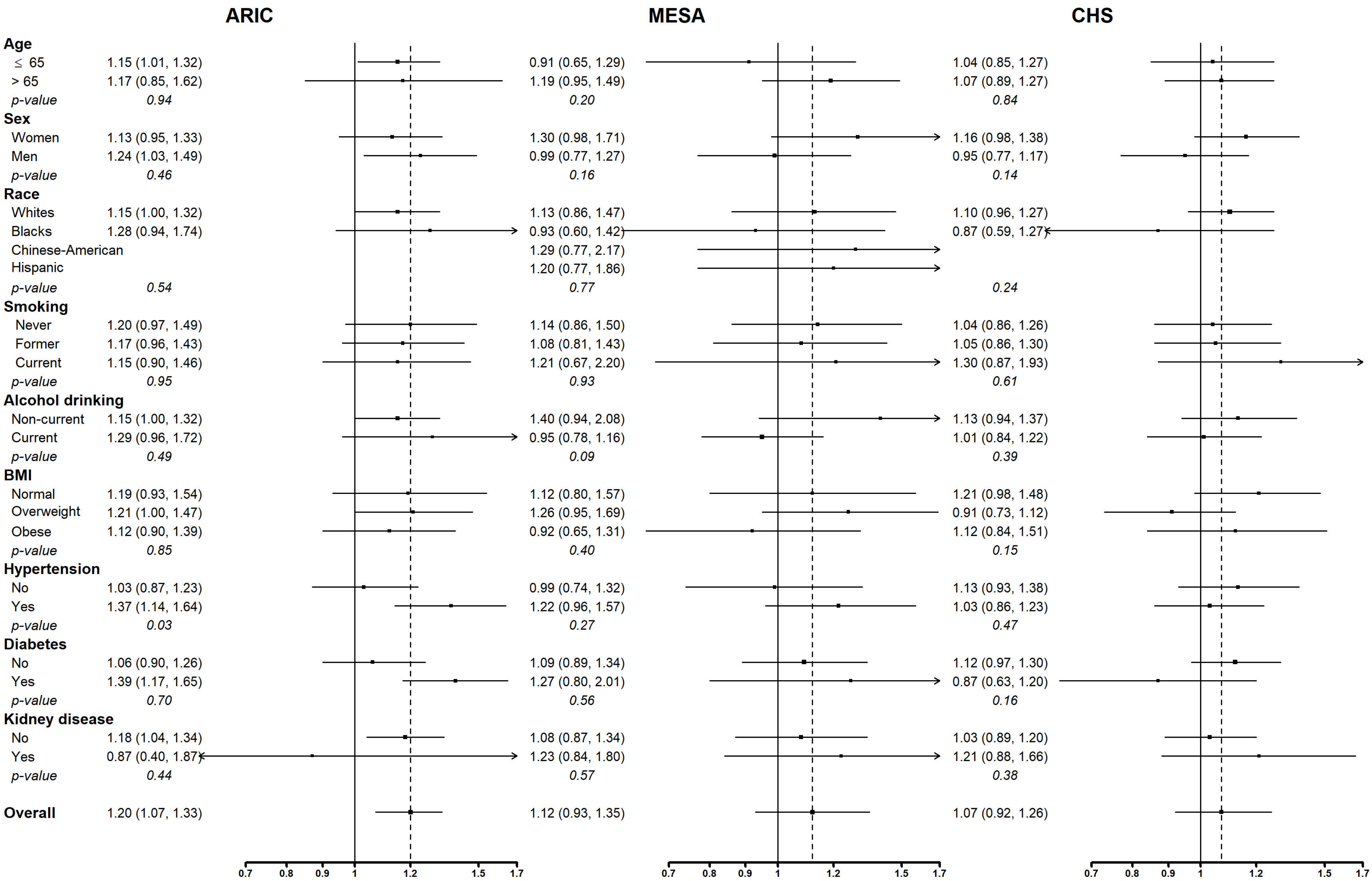
